# Trisomy 21 Impairs Development of Enteric Neural Crest-Derived Cells via SOD1-Mediated RET Dysregulation

**DOI:** 10.64898/2026.06.30.735397

**Authors:** Karan Singh, Frank Liu, Andy Zhao, Iman Lohraseb, Teresa Davoli

## Abstract

Hirschsprung disease (HSCR) is a rare congenital disorder of the enteric nervous system (ENS), marked by the absence of enteric ganglia along variable lengths of the distal gastrointestinal tract, resulting in functional intestinal obstruction. Individuals with Trisomy 21 (Down syndrome) face a 50- to 100-fold increased risk of HSCR relative to the general population, yet the molecular basis of this susceptibility remains poorly understood. Here, we investigated this association using isogenic induced pluripotent stem cells (iPSCs) derived from a mosaic individual with Down syndrome, enabling direct comparison of Trisomy 21 and Disomy 21 cells within an identical genetic background following differentiation into enteric neural crest-derived cells (ENCDCs). Trisomy 21 ENCDCs exhibited reduced proliferative and migratory capacity and an impaired ability to differentiate into enteric neurons relative to Disomy 21 controls. These phenotypes were accompanied by decreased *RET* expression at both the transcript and protein levels, together with broad downregulation of the *RET* gene regulatory network, including *GDNF*, *GFRA1*, *EDNRB*, *SEMA3C*, and *NRG1*, and of cell cycle and DNA replication pathways. Strikingly, we identified *SOD1*, a chromosome 21-encoded antioxidant enzyme not previously linked to RET regulation, as a dosage-sensitive driver of this effect: SOD1 overexpression in disomic ENCDCs was sufficient to suppress RET, whereas shRNA-mediated knockdown of *SOD1* in Trisomy 21 ENCDCs restored *RET* expression. Mechanistically, Trisomy 21 ENCDCs displayed markedly elevated catalase and a redox imbalance, and exogenous hydrogen peroxide recapitulated *RET* suppression in disomic cells, implicating oxidative stress as a mediator of RET downregulation. Collectively, these findings establish Trisomy 21 dosage effects as disruptors of RET-dependent enteric neural crest development and implicate SOD1-driven oxidative stress as a candidate mechanism, providing a framework for understanding the elevated risk of HSCR in Down syndrome.

## INTRODUCTION

Hirschsprung disease is a rare neurodevelopmental disorder marked by the partial or total absence of enteric ganglia along varying lengths of the distal gastrointestinal tract, resulting in severe intestinal dysmotility and functional intestinal obstruction. It arises when enteric neural crest-derived cells (ENCDCs) fail to adequately proliferate, migrate, survive, and differentiate during embryonic colonization of the gut, leaving the affected segments aganglionic, [1], [2], [3]. Its incidence is approximately 1 in 3,500–5,000 live births, but rises dramatically in Down syndrome (DS) to roughly 1 in 40–50, a 50- to 100-fold elevation that makes trisomy 21 the clearest syndromic setting in which to study ENS vulnerability [4], [5], [6].

The receptor tyrosine kinase RET is a cell-surface transmembrane receptor that forms a signaling complex with GDNF and the co-receptor GFRA1 to regulate enteric neural crest-derived cell (ENCDC) survival, proliferation, migration, differentiation, and neuronal maintenance. Reduced *Ret* gene dosage in mice is sufficient to cause colonic aganglionosis, underscoring the critical role of the GDNF–GFRA1–RET signaling axis in enteric nervous system development, [7], [8], [9]. A second axis, EDNRB and endothelin-3, also contributes to ENS patterning, and RET and EDNRB are now understood as interacting components of a broader gene regulatory network that shapes HSCR susceptibility [1], [10]. Reduced RET expression is likely to increase susceptibility to Hirschsprung disease; however, additional genetic or environmental modifiers are probably required for the development of overt aganglionosis [11], [12].

A striking genetic observation points to the *RET* gene, which is located on chromosome 10q11.2, as a target of chromosome 21 dosage. The common *RET* enhancer variant rs2435357 (C>T in intron 1) lowers *RET* expression and is the strongest common risk factor for HSCR, with the risk T allele enriched in HSCR⁺DS⁻ patients (allele frequency ∼0.61). However, DS⁺HSCR⁺ individuals carry the T risk allele at only an intermediate frequency (∼0.41)—lower than that in HSCR⁺DS⁻ individuals (∼0.61) and higher than that in individuals with DS without HSCR or HSCR⁻DS⁻ controls (∼0.26).[6]. DS individuals, therefore, require less of the common *RET* risk allele to cross the disease threshold, demonstrating an interaction between chromosome 21 dosage and RET-dependent susceptibility and implying that trisomy 21 itself supplies part of the burden on *RET*, most plausibly through increased dosage of chromosome 21 genes that suppress it [6]. Several chromosome 21-linked modifiers fit this model. A dose-sensitive chromosome 21 scan implicated *DSCAM* in both DS-associated and non-syndromic HSCR [13]. *COL6A1* and *COL6A2*, which encode collagen VI subunits on chromosome 21, provide a strong mechanistic link, as excess collagen VI production slows enteric neural crest migration and is enriched around ganglia in HSCR tissue, especially in DS [14], [15].

More recent work nominates *SOD1*, a chromosome 21-encoded cytoplasmic enzyme, as an additional candidate effector, showing that trisomy-level *Sod1* can suppress neuronal *Ret* and shift ENS cell states toward glial and progenitor programs [16]. Beyond frank aganglionosis, Trisomy 21 perturbs ENS structure more broadly. In Ts65Dn and Tc1 mouse models, early neural crest migration is largely preserved, yet submucosal neuron density is reduced and colonic motility impaired, indicating a wider spectrum of ENS abnormalities than distal aganglionosis alone [5]. Together these findings support a threshold model in which Trisomy 21 raises HSCR risk by converging on RET-centered pathways through extracellular matrix changes, chromosome 21 susceptibility loci, and candidate suppressors of neurogenesis [14]. A fundamental question remains: is the elevated HSCR risk in DS driven exclusively by chromosome 21 dosage acting cell-intrinsically, or does it require interaction with modifiers elsewhere in the genome? Human pluripotent stem cell systems that generate vagal enteric neural crest progenitors and enteric neurons allow these dosage effects to be tested directly in human cells [17].

Here, we study how trisomy 21 affects the differentiation of human enteric neural crest-derived cells and enteric neurons, using isogenic induced pluripotent stem cell lines from a single mosaic DS donor that differ only in the presence of a supernumerary chromosome 21 [18]. We find that trisomy 21 ENCDCs show reduced *RET* expression with impaired proliferation and migration, and we identify *SOD1* as a candidate chromosome 21 effector whose elevated dosage suppresses *RET*, providing a mechanistic framework for the elevated risk of HSCR in Down syndrome.

## RESULTS

### Trisomy 21 induces a defect in enteric neural crest derived cells differentiation

To study the effect of chromosome 21 Trisomy on enteric neural crest cells differentiation, we used isogenic iPSCs that are either disomic or trisomic for chromosome 21. These cells were derived from an individual with mosaic Down syndrome and obtained from the WiCell Research Institute, USA [18]. We first validated by whole-genome sequencing that the two trisomic clones indeed harbored a gain of chromosome 21, whereas the disomic clone did not.

To assess the effect of chromosome 21 dosage on enteric neural crest cell development, isogenic Disomy 21 and Trisomy 21 iPSCs were subjected to a staged induction protocol, transitioning through Day 0–2, Day 2–6, and Day 6–14 induction media. By Day 14, cells had acquired an enteric neural crest-derived cells (ENCDCs) identity, as confirmed by expression of the early lineage marker RET (Fig. 2c), and were subsequently replated in enteric neural differentiation medium to generate enteric neurons by Day 45. The protocol was validated by the expected expression dynamics: stem cell markers such as *OCT4* decreased during differentiation, while regional differentiation markers such as *HOXB3* increased (Fig. 2b). Additionally, Fig. 2a-c shows data from the control iPSC line, confirming that the differentiation protocol is effective, as previously validated in the WA01 line (WiCell Research Institute) (Supplementary Fig. 1f).

Next, we reseeded day 12 ENCDCs and assessed their proliferative capacity. Quantitative analysis revealed that Trisomy 21 ENCDCs exhibited a statistically significant increase in population doubling time compared to isogenic Disomy 21 controls (specifically, +3 hours for Trisomy 21-1 and +14 hours for Trisomy 21-2) indicative of a reduced proliferative rate in trisomic ENCDCs (Fig. 1c-d). Interestingly a similar analysis performed on iPSCs showed no significant difference in proliferative capacity between Trisomy and Disomy 21 cells (Supplementary Fig.1d-e).

**Figure 1.**
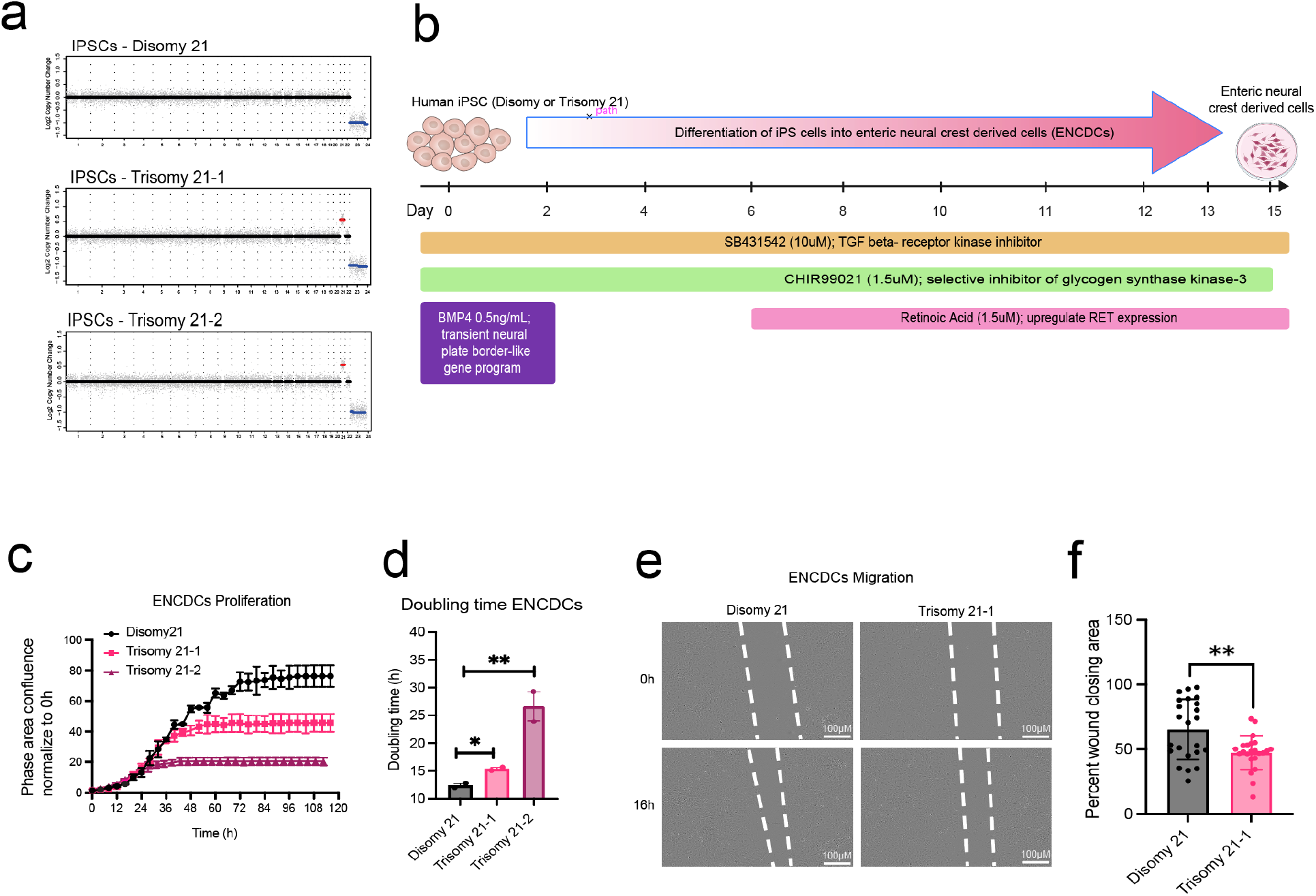
Reduced Proliferative and Migratory Capacity of iPSC-Derived Trisomy 21 ENCDCs. **a,** Chromosomal copy number confirmation by low-pass whole-genome sequencing (0.5×) performed on Disomy 21 (D21), Trisomy 21-1 (T21-1), and Trisomy 21-2 (T21-2) iPSC lines. **b,** Schematic of the ENCC derivation protocol from iPSCs. **c,** Cell proliferation assay of ENCDCs derived from Disomy 21 and Trisomy 21 iPSC lines. Cells were seeded at a density of 150,000 cells per well in 12-well plates on Day 11 of differentiation, and proliferation was monitored by live-cell imaging using the Incucyte system. Representative data from one of three independent experiments are shown. **d,** Quantification of doubling time from cell proliferation using an R script (further details are provided in the Methods section). **e,** ENCDCs migration assay performed at Day 14 of differentiation. Representative images are shown from one of 23 fields acquired per group. **f,** Quantification of wound closure area in ENCDCs following *in vitro* scratch assay, assessed at 16 hours post-scratch. Wound area was analyzed using Incucyte S3 (Sartorius) live-cell imaging system. Data are presented as percentage wound closure normalized to Disomy 21 controls. *n* = 23 images per group. Statistical significance was examined by nonparametric Welch’s t test comparisons test (Fig 1d) or non-parametric Mann-Whitney U test (Fig. 1f). Data are represented as mean ± SEM. p < 0.05, p < 0.01, p < 0.001, p < 0.0001; ns, not significant.

To further investigate the effect of Trisomy 21 on other components of enteric nervous system development, we evaluated the migratory capacity of ENCDCs using an *in vitro* scratch assay. Trisomy 21-1 ENCDCs exhibited a significant reduction in wound closure compared to isogenic Disomy 21 controls at 16 hours post-scratch (72.6 ± 4.21% vs. 100.0 ± 7.41%; mean ± SEM), representing a ∼27.4% decrease in migratory capacity (Fig. 1e–f).

To determine whether Trisomy 21 affects the neurogenic differentiation potential of iPSC-derived ENCDCs, we examined their capacity to differentiate into enteric neurons. Trisomy 21 ENCDCs demonstrated a significantly compromised neuronal differentiation capacity compared to isogenic Disomy 21 controls. Immunofluorescence analysis revealed a significant reduction in the expression of the pan-neuronal marker TUJ1 in both Trisomy 21-1 and Trisomy 21-2 ENCDCs compared to Disomy 21 controls, specifically, a ∼90% (p=0.005) and ∼80% reduction (p= 0.05) in Trisomy 21-1 and Trisomy 21-2, respectively (Supplementary Fig. 1b–c). Altogether, these data suggest that ENCDCs with trisomy 21 are impaired in their proliferative capacity, migratory capacity, and ability to differentiate into enteric neurons.

### RET expression is impaired in Trisomy 21 cells during enteric neuron differentiation

Overall, the phenotypic data indicate that Trisomy 21 lines have a reduced capacity to differentiate into ENCDCs (Fig. 1). To investigate the molecular basis of this phenotype, we analyzed lineage-specific gene expression by qRT-PCR during differentiation of isogenic iPSCs into ENCDCs. As differentiation progressed, both disomic and Trisomy 21 iPSCs exhibited an approximately 95% reduction in the pluripotency marker OCT4 and a 4–6-fold increase in the region-specific marker HOXB3 at day 15 relative to day 0, confirming successful neural crest induction in both cell lines (Fig. 2a–c). We next turned to the Hirschsprung disease–associated gene *RET*, a marker of ENCDCs, which is normally upregulated during ENCDCs differentiation and is necessary for their function and differentiation capacity [2]. We found that *RET* was significantly upregulated during disomic 21 ENCDCs differentiation from iPSCs, with increases of approximately 9.4-fold and 4.4-fold from day 0 to days 15 and 20 post-differentiation, respectively (Fig. 2a–c). In contrast, both Trisomy 21-1 and Trisomy 21-2 derived ENCDCs exhibited an approximately 62% reduction in *RET* expression compared with disomic cells at day 15 post-differentiation (*p* < 0.05). By day 20, *RET* transcript levels were reduced by 69% and 87% in Trisomy 21-1 and Trisomy 21-2 cells, respectively (Fig. 2c). Consistent with the transcriptional data, RET protein expression was also significantly reduced in Trisomy 21-derived ENCDCs on day 15 post-differentiation, with decreases of 74% (p = 0.005) and 68% (p = 0.05) in Trisomy 21-1 and Trisomy 21-2 cells, respectively, as determined by immunofluorescence (Fig. 2d–e). Together, these data demonstrate that Trisomy 21 reduces RET expression at both the transcript and protein levels during ENCDCs differentiation, identifying impaired RET regulation as a likely molecular contributor to the differentiation and proliferation defects observed in Trisomy 21 ENCDCs.

**Figure 2.**
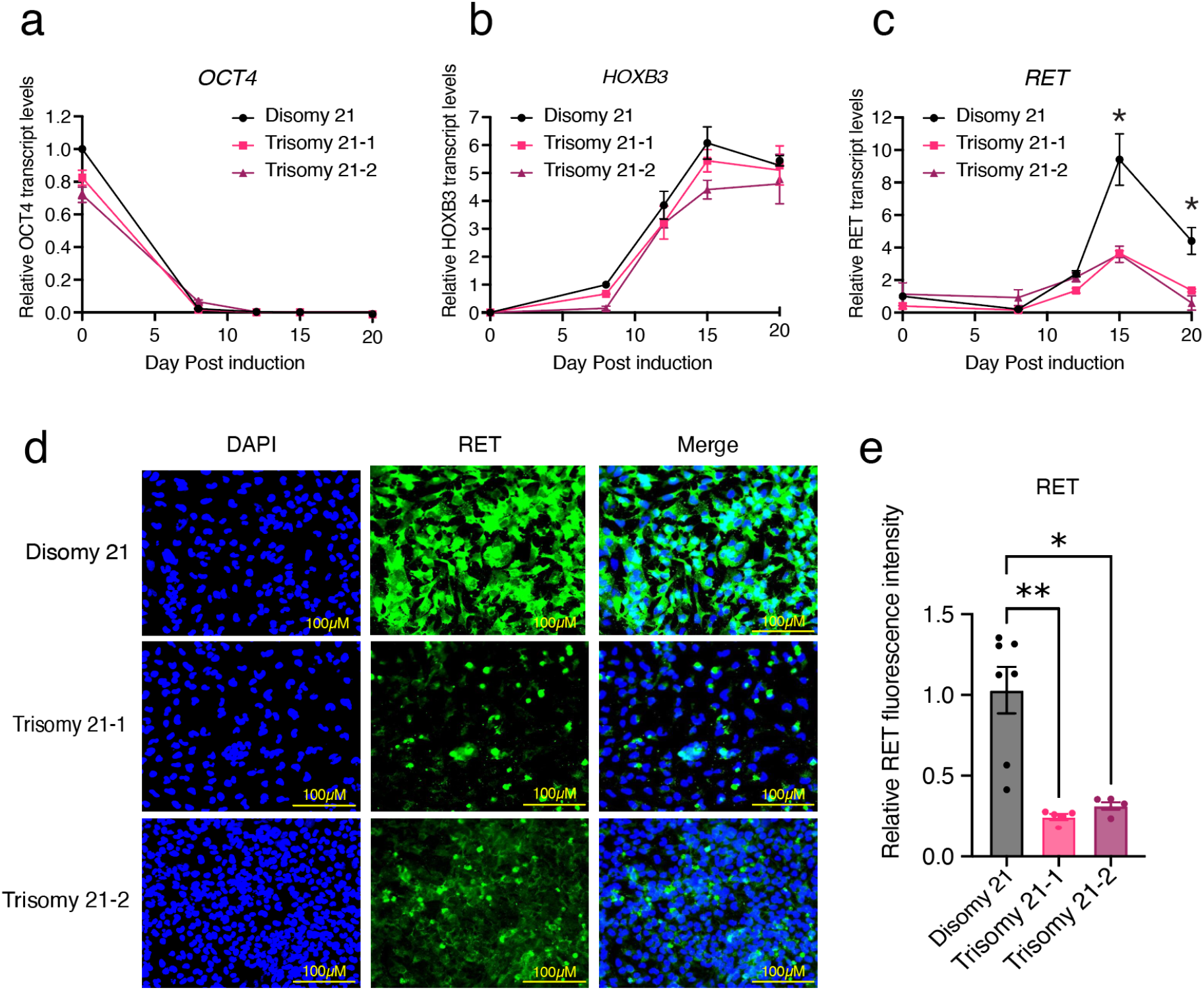
RET expression is impaired in Trisomy 21. **a–c,** Temporal gene expression profiling during iPSC-to-ENCDC differentiation. Relative transcript levels of *OCT4*, *HOXB3*, and *RET* were measured at sequential differentiation time points (*n* = 2 biological and technical replicates for each genotype; experiment independently repeated twice). **d,** Immunofluorescence staining for RET in Disomy 21, Trisomy 21-1, and Trisomy 21-2 ENCDC cells at Day 15 of differentiation. Cells were fixed with 2% paraformaldehyde (PFA) and stained with an anti-RET antibody (green); nuclei were counterstained with DAPI (blue). Scale bars, 100 µm. Representative images from one experiment are shown. **e,** Quantification of RET expression per cell, calculated as total fluorescence intensity normalized to total nuclei count. Statistical significance of Fig. 2c, and 2e was assessed using nonparametric tests, including the Kruskal–Walli’s test with Dunn’s multiple comparisons test. Data are presented as mean ± SEM. p < 0.05, p < 0.01, p < 0.001, p < 0.0001; ns, not significant.

### Trisomy 21 broadly downregulates Hirschsprung disease–associated genes and cell cycle pathways

To assess in an unbiased manner the gene regulatory changes driven by trisomy 21, we performed bulk RNA sequencing on isogenic Disomy 21 and Trisomy 21 cells at sequential stages of differentiation from iPSCs toward ENCDCs, specifically at day 0, day 12, and day 15 post-induction. Differential expression analysis of Trisomy 21 versus Disomy 21 ENCDCs revealed that numerous genes were significantly dysregulated, with 693 up-regulated genes and 439 down-regulated genes on Day 12 (FDR < 0.05) (Fig. 3a). Notably, the majority of HSCR associated genes were significantly downregulated (FDR < 0.05) in Trisomy 21 relative to Disomy 21 cells, including *RET*, *GDNF*, *GFRA1*, *EDNRB*, *SEMA3C*, *NRG1*, *SOX10*, *SLC27A4*, and *TCF4* (Fig.3b,c,g). This downregulation was further confirmed at the level of individual transcript read counts for representative genes, including *RET*, *GFRA1*, *GDNF*, and *EDNRB* (Fig. 3c). Together, these data suggest that Trisomy 21 broadly suppresses the *RET* gene regulatory network in enteric neural crest-derived cells, including *RET*, its ligand *GDNF*, its co-receptor *GFRA1*, and additional Hirschsprung disease (HSCR)-associated genes, such as *EDNRB*, *SEMA3C*, and *NRG1, TCF4*.

**Figure 3.**
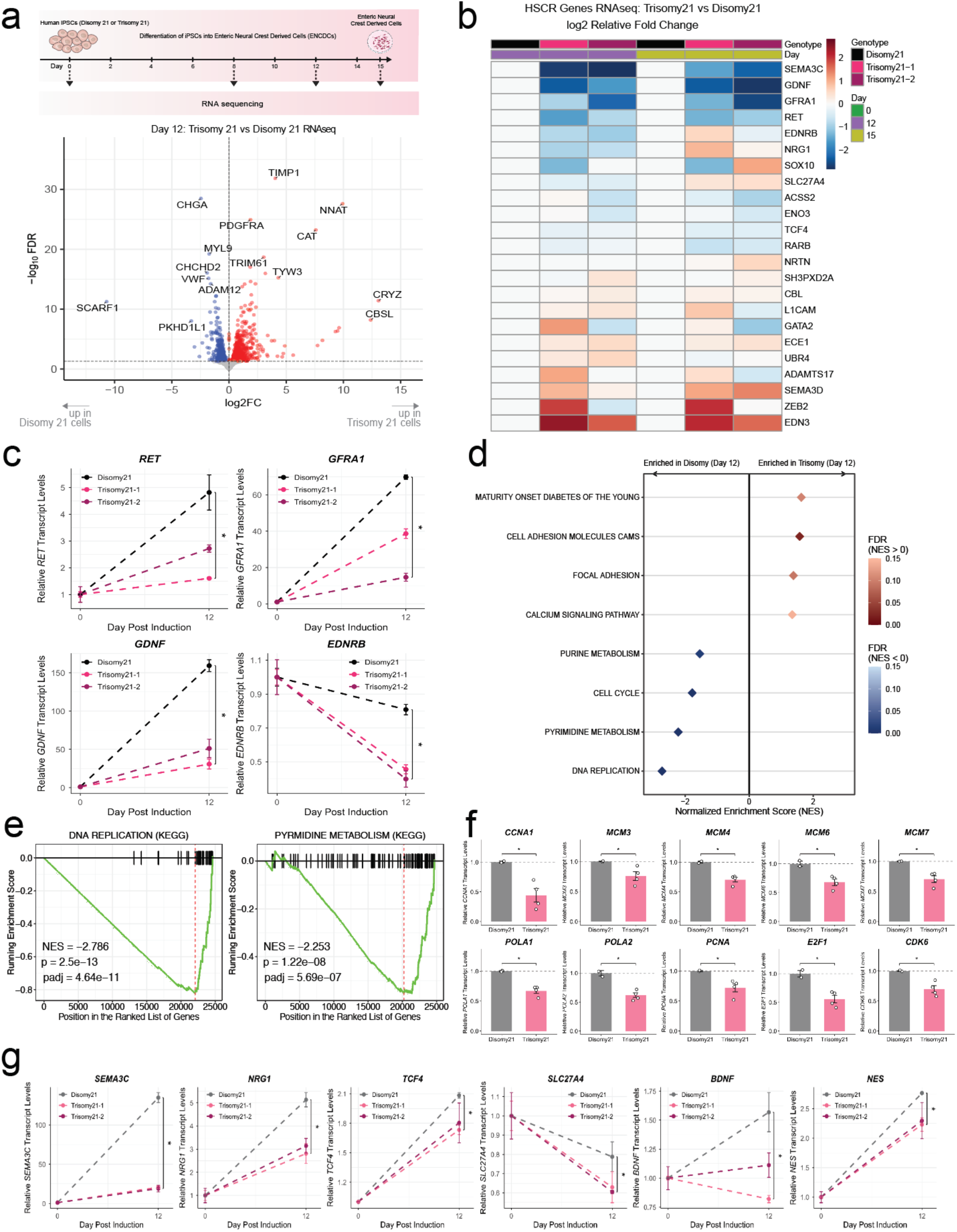
Trisomy 21 ENCDCs exhibit reduced expression of Hirschsprung disease-associated genes. **a,** Schematic of the ENCDCs derivation protocol from iPSCs, along with volcano plot illustrating Trisomy 21 vs Disomy 21 significantly differentially expressed genes on day 12 post induction. **b,** Heatmap of Trisomy 21 vs Disomy 21 log2 relative fold change. TPM values were averaged across technical replicates and normalized on days 12 and 15 with respect to day 0 expression levels within each genotype. Next, a second normalization step was performed whereby fold changes observed across time in the Trisomy 21 clones were normalized with respect to the observed fold change across time in the Disomy 21 clone. Finally, log2 was applied on top of all the relative fold change values, producing the heatmap shown. **c,** Relative transcript levels of genes in the *RET* regulatory network (*RET*, *GFRA1*, *GDNF*, *EDNRB*) on Day 12 of iPSC differentiation into ENCDCs. P-values were computed using one-sided Wilcoxon rank-sum test (* indicates p=0.0667, the lowest possible p value achievable from the sample size). **d,** KEGG gene set enrichment analysis (GSEA) of differentially expressed genes in Trisomy 21 versus Disomy 21 ENCDCs on Day 12. **e,** GSEA enrichment plots for KEGG DNA REPLICATION and KEGG PYRMIDINE METABOLISM pathways. **f,** Relative expression of cell cycle- and DNA replication-associated genes, including *CCNA1*, *MCM3*, *MCM4*, *MCM6*, *MCM7*, *POLA1*, *POLA2*, *PCNA*, *E2F1*, and *CDK6*, in Day 12 Disomy 21 and Trisomy 21 ENCDCs. **g,** Relative transcript levels of other Hirschsprung-related genes (*SEMA3C*, *NRG1*, *TCF4*, *SLC27A4*, *BDNF*, *NES*) on Day 12 of iPSC differentiation into ENCDCs. P-values were computed using one-sided Wilcoxon rank-sum test (* indicates p=0.0667, the lowest possible p value achievable from the sample size).

To identify the broader pathways affected by Trisomy 21, we next performed gene set enrichment analysis (GSEA) of differentially expressed genes in Trisomy 21 versus Disomy 21 ENCDCs. Strikingly, this analysis revealed significant downregulation of pathways associated with cell cycle progression and proliferation, including purine metabolism (NES = -1.551, padj = 0.014), cell cycle (NES = -1.795, padj = 8.98e-4), pyrimidine metabolism (NES = −2.205, padj = 7.4e−07), and DNA replication (NES = −2.735, padj = 4.83e−11) (Fig. 3d, e). Consistent with this, individual cell cycle associated genes, including cyclins (CCNA1), minichromosomal maintenance genes (*MCM3*, *MCM4*, *MCM6*, *MCM7*), DNA polymerase subunits (*POLA1*, *POLA2*), *PCNA*, *E2F1*, and CDK6, were significantly downregulated in Trisomy 21 relative to Disomy 21 cells (Fig. 3f). Collectively, these findings indicate that Trisomy 21 impairs the expression of both HSCR associated genes (Supplementary Fig. S2C and core proliferative machinery required for proper enteric nervous system development. Collectively, these findings indicate that Trisomy 21 perturbs the expression of critical genes and signaling pathways required for proper enteric nervous system development and suggest that the resulting impairment in proliferative capacity and altered nucleotide metabolism may contribute to the developmental defects observed in Trisomy 21-derived ENCDCs.

### SOD1 dosage regulates RET expression in ENCDCs

To identify which chromosome 21 gene(s) might drive the reduction in RET expression in ENCDCs, we first examined which genes on chromosome 21 are significantly upregulated in Trisomy 21 versus Disomy 21 cells across different stages of differentiation. Among all 153 expressed chromosome 21 genes, 45 were significantly upregulated at day 12 (FDR < 0.05) and 63 were upregulated on day 0 iPSCs (FDR < 0.05) based on our RNA-seq analysis (Fig. S3c). Among these candidates was superoxide dismutase 1 (*SOD1*), located at cytoband 21q22.11. Consistent with a gene dosage effect, *SOD1* was upregulated in Trisomy 21 relative to Disomy 21 cells, reaching significantly higher levels at day 8 post-differentiation (Fig. 4a). This expression pattern suggested that elevated *SOD1* may contribute to the impaired differentiation of Trisomy 21 cells into ENCDCs and to the reduced *RET* expression observed in these cells.

**Figure 4.**
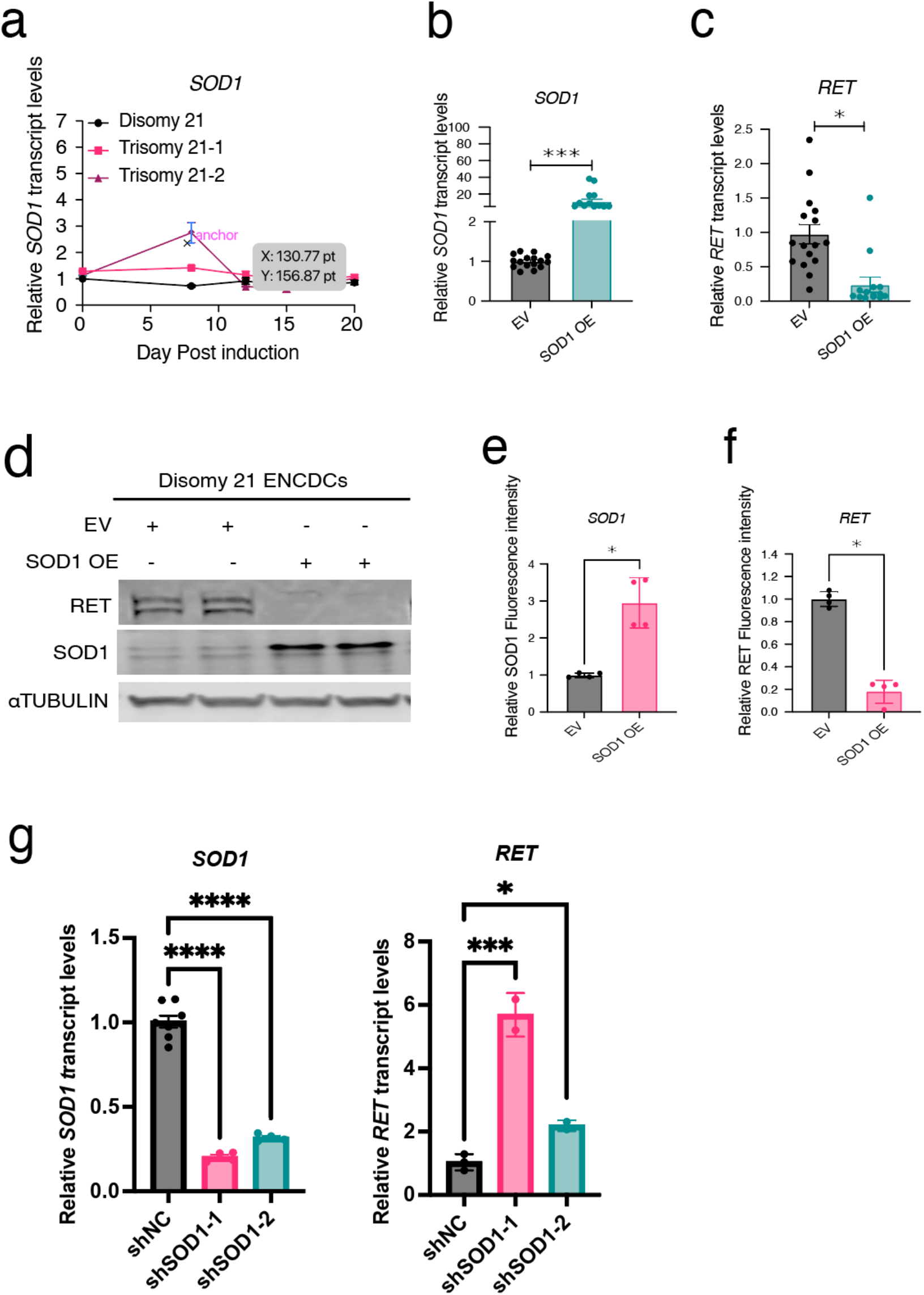
SOD1 overexpression modulates RET expression in ENCDCs. **a,** Relative transcript levels of *SOD1* at Day 14 of differentiation in Disomy 21 iPSCs overexpressing *SOD1* or carrying an empty vector control (*n* = 4; two technical replicates per sample). **b,c,** Relative transcript levels of *RET* and *SOD1* in *SOD1*-overexpressing ENCDCs at Day 14 of differentiation (*n* = 3 independent experiments; 2–3 technical replicates per sample). **d,** Representative immunoblots from one of two independent experiments showing RET and SOD1 protein levels in ENCDCs lysates collected at Day 14 from *SOD1*-overexpressing and empty vector control conditions. **e,f,** Quantification of RET and SOD1 immunoblot signal intensities normalized to α-TUBULIN. Data are shown from two independent experiments (*n* = 4; two technical replicates per group). **g**, Relative *SOD1* and *RET* transcript levels in Trisomy 21-1 ENCDCs expressing shRNA against *SOD1* or a negative control shRNA (shNC) at Day 14 of differentiation (*n* = 2-4 biological replicates; two technical replicates per group). Statistical significance for Fig. 4b,c and Fig. 4e,f was assessed using the nonparametric Mann– Whitney U test. Statistical significance for Fig. 4g was assessed using the nonparametric Kruskal– Walli’s test with Dunn’s multiple comparisons test. Data are presented as mean ± SEM. p < 0.05, p < 0.01, p < 0.001, p < 0.0001; ns, not significant.

To test directly whether increased *SOD1* dosage affects *RET* expression, we lentiviral transduced Disomy 21 iPSCs with a vector expressing human *SOD1* and measured *RET* expression following differentiation into ENCDCs. Increased *SOD1* dosage led to a significant decrease in RET expression at both the transcript and protein levels (Figure 4b-f). We next performed the complementary experiment, depleting *SOD1* by shRNA-mediated transduction of Trisomy 21 cells. Knockdown of *SOD1* resulted in a significant increase in RET expression in Trisomy 21 ENCDCs (Fig.4g), supporting the conclusion that *SOD1* gene dosage plays an important role in regulating RET at both the transcriptional and protein levels. Collectively, these findings identify *SOD1* as a chromosome 21 dosage-sensitive gene that negatively regulates RET expression in enteric neural crest-derived cells, nominating *SOD1* overexpression as a candidate mechanism linking Trisomy 21 to *RET* suppression and the elevated risk of Hirschsprung disease in Down syndrome.

### SOD1 regulates RET expression through an oxidative stress-dependent mechanism

Given the established role of SOD1 in regulating cellular redox homeostasis, we next examined whether oxidative stress might mediate the effect of elevated SOD1 on RET expression. We first tested whether SOD1 might act through a nuclear, potentially transcriptional, role, as has been recently reported [19]. We performed nuclear and cytoplasmic fractionation in Disomy 21 ENCDCs and in ENCDCs overexpressing SOD1. SOD1 was not detectable in the nuclear fraction, with the vast majority of the signal localized to the cytoplasm (Fig. 5a), arguing against a direct nuclear or transcriptional role for SOD1 in RET regulation.

**Figure 5.**
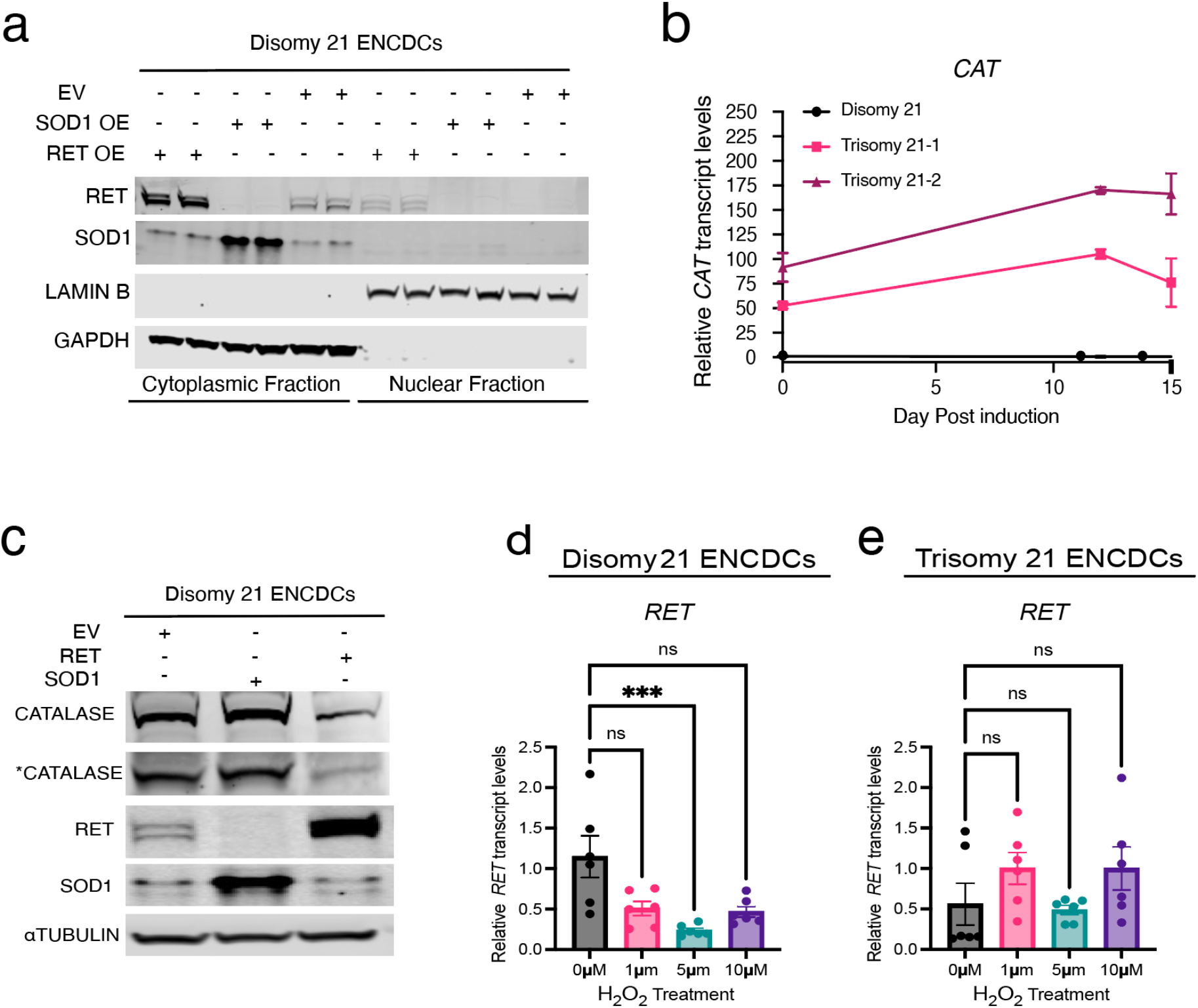
SOD1 regulates RET expression through an oxidative stress-dependent mechanism. **a,** SOD1 does not localize to the nucleus. Representative immunoblots from one experiment showing RET and SOD1 protein levels in ENCDC lysates collected at Day 14 from RET- or SOD1-overexpressing and empty vector control conditions. Data are shown from one experiment (*n* = 2 technical replicates per condition). **b,** Relative *CAT* transcript levels during iPSC differentiation in Disomy 21, Trisomy 21-1, and Trisomy 21-2 lines (*n* = 2 replicates per group) are shown. Data are representative of one of two independent immunoblots. **C,** Representative immunoblots from one of two independent immunoblots showing RET, CAT (Catalase) and SOD1 protein levels in ENCDCs lysates collected at Day 14 from SOD1 *or* RET - overexpressing and empty vector control conditions. Data are shown from one biological replicate with two technical replicates per group (*n* = 1 biological replicate; 2 technical replicates per group). Data are representative of one of two independent immunoblot experiments. **d, e,** *RET* transcript levels in Disomy 21 and Trisomy 21-1 ENCDCs treated with doses of 0 µM, 1 µM, 5 µM, or 10 µM H₂O₂ for 4 hours on Day 15 post-differentiation were measured by qRT-PCR as described in the Methods. qRT-PCR values were normalized to *Gapdh* and plotted relative to Disomy 21 0 µM. (*n* = 3 biological replicates with two technical replicates per treatment condition). Data are representative of one experiment. Statistical significance of Fig. 5d, and 5e was assessed using nonparametric tests, including the Kruskal–Walli’s test with Dunn’s multiple comparisons test. Data are presented as mean ± SEM. p < 0.05, p < 0.01, p < 0.001, p < 0.0001; ns, not significant.

Given the role of SOD1 in oxidative stress, we next examined the level of catalase, a key antioxidant enzyme, in trisomic and disomic cells. Catalase expression was markedly elevated in Trisomy 21 relative to Disomy 21 cells, by approximately 50 to 100-fold (Fig. 5b), suggesting that trisomic cells experience substantial oxidative stress and upregulate catalase to withstand this damage. Consistent with a coordinated redox network, SOD1 overexpression increased catalase levels, whereas RET overexpression decreased them (Fig. 5c), pointing to a reciprocal co-regulation among SOD1, catalase, and RET.

Finally, reasoning that H₂O₂ generated downstream of elevated SOD1 might mediate RET suppression, we exposed Disomy 21 ENCDCs to increasing concentrations of H₂O₂ to mimic elevated oxidative stress. H₂O₂ treatment resulted in a dose-dependent reduction in *RET* expression (Fig. 5d). Importantly, this effect was not observed in Trisomy 21 cells (Fig. 5e), indicating that the response is specific to disomic cells. Collectively, these findings suggest that elevated *SOD1* expression contributes to *RET* downregulation in Trisomy 21 ENCDCs, potentially through an oxidative stress-mediated mechanism.

## DISCUSSION

In the present study, we provide direct evidence that RET expression is significantly reduced in Trisomy 21 iPSC-derived ENCDCs, thereby identifying impaired RET signaling as a plausible pathogenic mechanism underlying Down syndrome-associated HSCR. Our findings are further supported by human in vivo datasets. In addition to reduced circulating RET protein levels [20], whole-blood RNA-sequencing data from the Human Trisome Project [21] indicate that RET expression is decreased in individuals with Trisomy 21. In the context of the well-established causal role of RET loss-of-function in HSCR [22], these observations reinforce the clinical relevance of our in vitro data and support the hypothesis that reduced RET expression is a contributing factor to the increased incidence of HSCR in Down syndrome. In line with the established functions of RET in regulating enteric neural crest cell proliferation, migration, and survival [2], [23], Trisomy 21 ENCDCs exhibited marked defects in both proliferative and migratory capacity relative to their isogenic disomic counterparts. Because coordinated migration and expansion of enteric neural crest cells are essential for complete colonization of the developing gastrointestinal tract [24], these defects provide a biologically coherent explanation for how reduced RET expression may predispose to aganglionosis. Together, these findings support a model in which RET insufficiency constitutes a mechanistic link between Trisomy 21 and heightened HSCR susceptibility.

Importantly, the transcriptional consequences of trisomy 21 in ENCDCs extend beyond RET alone. We observed significant downregulation of multiple genes of the *RET* regulatory net-work, including *EDNRB, GFRA1*, and *GDNF*, as well as several additional genes implicated in HSCR pathogenesis, including *TCF4, SEMA3C, NRG1*, *SLC27A4, BDNF,* and *NES*. These data suggest that trisomy 21 perturbs a broader *RET* gene regulatory network essential for enteric nervous system development rather than producing an isolated defect in RET expression. Such coordinated disruption would be expected to impair multiple cellular processes required for ENS formation, including lineage specification, migration, proliferation, and differentiation, thereby amplifying susceptibility to HSCR. Particularly notable is the reduction in *TCF4* transcript levels, a transcription factor with potential relevance to RET pathway regulation [25], raising the possibility that altered transcriptional control may further compound RET pathway dysfunction in Trisomy 21 ENCDCs.

An important unresolved question is whether RET downregulation in Trisomy 21 is driven predominantly by a single dosage-sensitive chromosome 21 gene or instead reflects the cumulative action of multiple dosage genes. Our data strongly implicate SOD1 as a major contributor. This interpretation is consistent with transcriptomic data from the Human Trisome Project [21] and with circulating proteomic data from studies by Sullivan et al., which showed reduced RET protein levels in individuals with Down syndrome [20] and which similarly demonstrate negative correlation of increased SOD1 expression accompanied by reduced RET expression in individuals with Down syndrome. Nevertheless, although our results establish SOD1 as sufficient to repress RET and necessary for maintaining reduced RET levels in Trisomy 21 ENCDCs, they do not exclude the possibility that additional chromosome 21 genes cooperate with SOD1 to shape the full Trisomy 21 transcriptional landscape.

A major advance of this study is the identification of SOD1 as a candidate dosage-sensitive regulator of RET expression in Trisomy 21 ENCDCs. SOD1, located on chromosome 21, is overexpressed in Down syndrome as a consequence of gene dosage imbalance. We found that ectopic SOD1 overexpression in disomic ENCDCs was sufficient to suppress RET expression, whereas shRNA-mediated SOD1 knockdown in Trisomy 21 ENCDCs restored RET levels. These complementary gain- and loss-of-function experiments provide strong evidence that increased SOD1 dosage contributes directly to RET repression in the trisomic context. Although SOD1 has classically been studied in the context of oxidative stress biology, our findings uncover a previously unappreciated role for SOD1 in the regulation of a key developmental signaling pathway required for enteric nervous system formation. To our knowledge, this represents the first in vitro demonstration that SOD1 dosage functionally modulates RET expression in enteric neural crest-derived cells.

Our findings are also concordant with emerging experimental evidence from model systems. A recent preprint by Grullon et al. (2026) reported that SOD1 trisomy is associated with reduced RET expression in the distal colon of postnatal day 0 mice and further suggested that an extra copy of SOD1 perturbs enteric nervous system development [17]. The mechanism by which SOD1 suppresses RET expression remains to be fully defined. Our preliminary data suggest that this regulation is unlikely to occur through direct transcriptional control by SOD1 itself and instead may be mediated by altered intracellular redox state. As a cytoplasmic antioxidant enzyme, SOD1 catalyzes the conversion of superoxide radicals to hydrogen peroxide (H₂O₂), thereby reshaping redox homeostasis. Consistent with this model, exogenous H₂O₂ treatment of disomic ENCDCs induced a dose-dependent reduction in RET expression, suggesting that elevated H₂O₂ or broader oxidative signaling may mediate the suppressive effect of increased SOD1 dosage on RET transcription.

A further consideration is that catalase is markedly overexpressed in trisomy 21 ENCDCs, reaching levels approximately 50- to 100-fold higher than in disomic cells. Notably, this dramatic induction was not recapitulated by SOD1 overexpression alone, indicating that elevated SOD1 dosage is not sufficient to drive such a pronounced increase in catalase expression. This observation suggests that the phenotypes we observe may reflect a combination of the gene-specific effect of SOD1 and a broader effect of whole-chromosome aneuploidy. Aneuploidy itself can elevate reactive oxygen species and oxidative stress; the convergence of this aneuploidy-driven oxidative burden with increased SOD1 expression would be expected to raise intracellular H₂O₂, which cells sense and counter by upregulating catalase activity. Despite this compensatory response, we hypothesize that Trisomy 21 cells sustain elevated H₂O₂, which in turn drives downstream events that reduce RET expression at both the transcript and protein levels. We therefore propose that a combination of the gene-specific effect of SOD1 and the whole-chromosome aneuploidy effect underlies the phenotypes observed in Trisomy 21 ENCDCs.

Taken together, our data support a model in which Trisomy 21 disrupts redox homeostasis through the convergence of two mechanisms: a gene-specific effect of elevated SOD1 dosage and a broader effect of whole-chromosome aneuploidy that itself increases oxidative stress. The resulting elevation in H₂O₂ suppresses RET expression and contributes to wider disruption of the developmental pathways required for enteric nervous system formation. These findings provide a mechanistic framework linking chromosome 21 dosage imbalance to impaired ENS development and offer a biologically grounded explanation for the increased susceptibility to HSCR in individuals with Down syndrome.

This study has several limitations. Most notably, the Trisomy 21 iPSCs used here were derived from a single individual with mosaic Down syndrome, whereas control lines were obtained from two independent individuals. Although the use of isogenic comparisons strengthens causal inference, validation across additional Trisomy 21 individual-derived iPSC lines will be important to establish the generalizability of these findings. Future studies incorporating a larger number of genetically independent lines differentiated into ENCDCs will enable more robust assessment of RET dysregulation and its upstream determinants. A second limitation is the absence of direct in vivo validation in human intestinal tissue. Analysis of single-cell RNA-sequencing datasets from Down syndrome colon specimens would provide an important opportunity to determine whether the transcriptional abnormalities observed in vitro are recapitulated in the native tissue context. However, such datasets remain difficult to obtain because of the scarcity of Down syndrome colon tissue and associated ethical and technical constraints. As these resources become available, they will be critical for validating the translational relevance of the present findings.

Overall, our results suggest that Trisomy 21 gene dosage affects RET expression in Trisomy 21 ENCDCs, potentially through a gene dosage effect of SOD1. These findings provide evidence that increased SOD1 dosage may contribute to RET dysregulation and, consequently, to impaired enteric neural crest cell development in Down syndrome.

## METHODS

### Induced Pluripotent Stem Cell Culture

Cytogenetically validated isogenic induced pluripotent stem cell (iPSC) lines, UWWC1-DS1 and UWWC1-DS4, derived from an individual with mosaic Trisomy 21, were obtained from the WiCell repository. These lines are hereafter referred to as Trisomy 21-1 and Trisomy 21-2, respectively. An isogenic disomic control line derived from the same individual with mosaic Trisomy 21, UWWC1-DS2U, was also obtained from WiCell and is hereafter referred to as Disomy 21. Establishment of these iPSC lines was previously described by Weick *et al.* (2013) [18]. Briefly, the parental fibroblasts were isolated from the thoracic skin of a one-year-old White male with mosaic Down syndrome, consisting of approximately 90% trisomic and 10% disomic cells. Reprogramming of these fibroblasts using the Yamanaka factors generated three stable iPSC clones: one disomic control line (DS2U) and two trisomic lines (DS1 and DS4), each retaining an additional copy of chromosome 21.

All iPSC lines were maintained on Cultrex-coated plates in mTeSR™ Plus medium (STEMCELL Technologies, Cat. No. 100-0276) with daily medium changes. For routine passaging, cultures were washed once with Versene (Thermo Fisher Scientific, 15040066) upon reaching approximately 30–40% confluence and then incubated with Accutase to obtain a single-cell suspension. Dissociated cells were collected by centrifugation at 150*g* for 5 minutes, resuspended in mTeSR™ Plus medium supplemented with 10 μM ROCK inhibitor Y-27632 (STEMCELL Technologies, 72302) to promote cell survival following passaging, and seeded onto freshly Cultrex-coated plates. The medium was replaced the following day and changed daily thereafter.

### Differentiation of iPSCs Into Enteric Neural Crest Derived Cells (ENCDCs)

To investigate the effect of chromosome 21 dosage on ENCDC differentiation, iPSCs were seeded at a density of 7.5 × 10⁵ cells per well in Cultrex-coated 6-well plates and expanded in mTeSR Plus medium until reaching approximately 80–90% confluence. Differentiation was initiated on Day 0 by transitioning cells to ENCDCs induction medium (Fig. 1b and Supplementary Table 1), which was maintained through Day 2. The medium was subsequently replaced with Day 2–6 induction medium, followed by a further transition to Day 6–14 induction medium on Day 6. By Day 14, cells had acquired an ENCDCs identity, as confirmed by expression of the early lineage markers SOX10 and RET. To generate enteric neurons, Day 14 ENCDCs were replated in enteric neural differentiation medium (Supplementary Table 2) and maintained until Day 60 with half-medium changes performed every other day.

### Genotyping of iPSC Lines

Genomic DNA was extracted from 1 × 10⁶ cells of each iPSC line (DS2U, DS1, and DS4). Sequencing libraries were prepared in-house, and low-pass whole-genome sequencing was performed at approximately ∼0.5X coverage. Copy number analysis and chromosomal dosage assessment were carried out using an in-house bioinformatic pipeline. All three lines were genotyped at the HSCR-associated variant rs2435357, a known regulatory variant in the *RET* proto-oncogene. Sanger sequencing confirmed that DS2U, DS1, and DS4 iPSCs were homozygous for the non-risk allele at this locus, excluding the RET risk haplotype (TT) as a confounding variable in subsequent differentiation experiments. Representative sequencing chromatograms and alignment data are provided in Supplementary Fig. S1a. Additionally, 30× whole-genome sequencing was performed on the iPSC lines used in this study. No pathogenic variants associated with Hirschsprung disease (HSCR) were identified. The sequencing data are available upon request.

### Overexpression of SOD1 and RET

To examine the effect of increased SOD1 or RET dosage on enteric neural crest-derived cell (ENCDC) differentiation, Disomy 21 iPSCs were seeded at a density of 7.5 × 10⁵ cells per well in Cultrex-coated 6-well plates and transduced with lentiviral particles carrying either a SOD1 overexpression construct (EF-1α intron A–SOD1), a RET overexpression construct (EF-1α intron A–RET), or an empty vector control. Three days after transduction, stable integrants were selected by culturing the cells in a medium supplemented with puromycin (0.5 µg/mL) for 3–5 days. Following puromycin selection, surviving cells were expanded and subsequently used for ENCDC differentiation.

ENCDCs differentiation was initiated once iPSCs reached approximately 80–90% confluence. On Day 0, cells were transferred to ENCDCs induction medium (composition provided in Supplementary and Fig S1b) and maintained in this formulation through Day 2. The medium was subsequently replaced with the corresponding stage-specific induction medium for Days 2–6, followed by a further transition to Day 6–14 induction medium to support continued differentiation and maturation. By Day 14, cells had acquired an ENCDCs identity, as confirmed by expression of the early lineage markers SOX10, HOXB3 and RET. Overexpression of each gene was independently validated at the transcript and protein levels by quantitative reverse transcription PCR (qRT-PCR) and western blotting, respectively. The effects of gene overexpression on the ENCDC differentiation trajectory were subsequently assessed.

### Knockdown of *SOD1* level

Trisomy 21-1 iPSCs were seeded at a density of 5 × 10⁵ cells per well in a cluster of 6-well plate. The following day, the culture medium was replaced and polybrene was added to a final concentration of 5 µg/mL to enhance transduction efficiency. Lentiviral particles were introduced at a multiplicity of infection (MOI) of 1–2 lentiviral particles per cell. Twenty-four hours post-transduction, the medium was replaced and cells were allowed to expand. On day 3 post-transduction, puromycin selection was initiated at a concentration of 0.5 µg/mL and maintained for 3–5 days to eliminate non-transduced cells. Following selection, surviving cells were re-seeded and subjected to directed differentiation as described above. Knockdown efficiency was assessed at day 14 post-differentiation.

### RNA Isolation, cDNA Synthesis, and Quantitative RT-PCR

We used the procedure described by Singh *et al.* (2022) [26]. Briefly, total RNA was extracted using QIAzol Lysis Reagent (QIAGEN, Cat. No. 79306) and purified using the RNeasy Mini Kit (QIAGEN, Cat. No. 74004) with on-column DNase digestion (QIAGEN, Cat. No. 79254) to eliminate genomic DNA contamination. One microgram of total RNA from each sample was reverse transcribed using the iScript cDNA Synthesis Kit (Bio-Rad, Cat. No. 1708891). Quantitative reverse transcription PCR (qRT-PCR) was performed using TaqMan Gene Expression Assays in combination with TaqMan Fast Universal PCR Master Mix (Thermo Fisher Scientific, Cat. No. 4367846) on an Applied Biosystems 7500 FAST Real-Time PCR System. Gene expression levels were normalized to the reference genes *B2M, GAPDH, or GUSB*, and relative expression was calculated using the 2^−^ΔΔCt^ method described in Singh et al., (2022) [27].

### RNA Sequencing

RNA was isolated as described above. RNA quantity and integrity were assessed at the NYU Genome Technology Center using an Agilent 2100 Bioanalyzer with RNA Nano Chips (Agilent, cat. no. 5067-1511). Only samples with RNA Integrity Number (RIN) values of 9.9–10 were included in downstream analyses to ensure high-quality input material. Library preparation was performed at the NYU Genomics Technology Center using 500 ng of total RNA per sample. Polyadenylated RNA was enriched via poly(A) selection, and stranded RNA-seq libraries were constructed using an automated library preparation workflow. Libraries were amplified for 11 PCR cycles and sequenced on an Illumina NovaSeq X platform using S1 100-cycle flow cells. Sequencing was performed in paired-end mode (50 bp reads) at a loading concentration of 12 pM with a 25% PhiX spike-in to ensure run quality.

### RNA Sequencing Data Analysis

Salmon was used to quantify raw transcript abundance counts from RNA sequencing fastq files. TPM expression values were calculated per transcript and averaged across technical replicate samples. DESeq2 was used to perform differential gene expression analysis amongst trisomy 21 vs disomy 21 samples. Volcano plots were produced using the Enhanced Volcano package in R. Gene set enrichment analysis (GSEA) was performed on top of the DESeq2 output, using the fgsea package in R with parameters minSize = 15, maxSize = 500, and nPerm = 10000. REACTOME and KEGG gene set libraries were used from the msigdbr package in R.

### Protein Extraction and Immunoblotting

We used the procedure described by Singh *et al.* [26]. Briefly, cells were lysed in RIPA buffer (50 mM Tris pH 7.5, 150 mM NaCl, 1% NP-40, 0.5% sodium deoxycholate, 0.1% SDS) supplemented with protease inhibitors (Roche Complete), phosphatase inhibitors (PhosSTOP), and 1 mM phenylmethylsulfonyl fluoride (PMSF; Cell Signaling). Lysates were clarified by centrifugation at 15,000 × g for 15 minutes at 4°C, and total protein concentration was determined using the Pierce BCA Protein Assay Kit (Thermo Scientific, cat. no. 23227). Equal protein amounts (25 µg per lane) were resolved on ExpressPlus 4–20% gradient SDS-PAGE gels (GenScript, cat. no. M42015) and subsequently transferred onto 0.2 µm nitrocellulose membranes (Bio-Rad, cat. no. 1620212) at 60 V for 2 hours. Membranes were blocked in Intercept PBS Blocking Buffer (LI-COR, cat. no. 927-70001) for 1 hour at room temperature and then incubated overnight at 4°C with the following primary antibodies: anti-RET (Thermo Scientific), anti-SOD1, anti-CAT and anti-α-TUBULIN or GAPDH, LAMIN B (all from Cell Signaling Technology). After washing, membranes were incubated with LI-COR IRDye fluorescent secondary antibodies (IRDye 800CW anti-rabbit and IRDye 680RD anti-mouse) for 1 hour at room temperature. Fluorescent signal detection and band quantification were performed using the LI-COR Odyssey CLx imaging system and Image Studio software (version 5.2).

### Immunocytochemistry

Immunostaining was performed following the protocol described in Single et al., 2022 [26]. Briefly, cells were seeded onto 8-chamber slides and fixed in 2% paraformaldehyde, followed by three washes with PBS. Non-specific binding was blocked using goat serum for 1 hour at room temperature. Slides were then incubated overnight at 4°C with primary antibodies directed against RET or TUJ1, diluted in 1X PBST, followed by three washes with PBST. Secondary antibody incubation was subsequently carried out in 1X PBST. Nuclear counterstaining was performed using DAPI at a concentration of 0.66 µg/µL for 5 minutes at room temperature, followed by an additional wash with PBST. Slides were mounted using an antifade reagent (Invitrogen ProLong Gold Antifade Mountant; P10144), allowed to cure at room temperature for 3 days, and subsequently sealed with colorless nail polish. Slides were stored at −20°C until image acquisition.

### Cell Migration Assay

To assess the migratory capacity of enteric neural crest-derived cells (ENCDCs), isogenic ENCDCs derived from iPSC lines with and without Trisomy 21 were subjected to a scratch-wound migration assay. On day 12 of differentiation, ENCDCs were seeded onto Cultrax-coated plates at a density of 5 × 10⁵ cells per well. Once the cells reached 100% confluency, a uniform scratch (wound) was manually created using a sterile pipette tip. A fresh differentiation medium was then added, and the plates were transferred to the Sartorius Incucyte S3 Live-Cell Imaging System for continuous monitoring. Images were acquired every 4 hours for 24 hours. Wound closure was subsequently quantified from the time-lapse images using an image analysis of two time point 0h or 16h post scratch or wound creation developed with Claude Sonnet 4.6.

### iPSCs and ENCDCs Proliferation Assays

To assess cell proliferation and determine doubling time, 1.5 × 10⁵ iPSCs (DS1, DS4, or DS2U) were seeded into each well of a 12-well plate. After 24 h, the culture medium was refreshed, and the plates were transferred to a Sartorius Incucyte S3 Live-Cell Imaging System for continuous monitoring over a 96 or 116h period. For each cell line, 3–4 replicate wells were imaged, with a minimum 16 fields acquired per well. Cell confluence data were analyzed using Sartorius Incucyte S3 Live-Cell Imaging System software to generate growth curves. An R-based script was used to identify the linear growth phase of each curve and calculate the corresponding growth rate (slope, m). Doubling time (DT) was calculated using the equation:

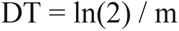

Only growth curves exhibiting a coefficient of determination (R²) ≥ 0.975 were included in the final analysis.

### Hydrogen Peroxide (H₂O₂) Assay

To assess whether H₂O₂ affects RET expression in enteric neural crest-derived cells (ENCDCs), isogenic ENCDCs derived from iPSC lines with and without Trisomy 21 were reseeded at a density of 5 × 10⁵ cells per well in Cultrex-coated 6-well plates on day 12 of differentiation. On day 15, cells were treated with increasing concentrations of H₂O₂ (0, 1, 5, or 10 μM) for 4 hours. Following treatment, total RNA was isolated, and cDNA was synthesized. RET expression was then quantified by quantitative reverse transcription PCR (qRT-PCR) as described above.

## AUTHOR CONTRIBUTIONS

**KS:** Conceptualization, Formal analysis, Investigation, Methodology, Project administration, Validation, Visualization, Writing – original draft, Writing – review & editing. **FL:** Formal analysis and Visualization. **AZ and IL:** Investigation. **TD:** Funding acquisition, Resources acquisition, Conceptualization, Visualization, Formal analysis, Supervision, Writing – review & editing.

## ACKNOWLEDGEMENTS

This work was supported by grants from the National Institutes of Health (R37 R37CA248631, R01 R01HG012590, and R01 R01DK135089), the Pershing Square Sohn Prize for Young Investigators in Cancer Research and a collaborative grant from the NFCR. We apologize to scientists in the field whose work we were unable to cite due to space constraints. We thank Princi Labana for analysis of the cell migration data. We thank Trinity de León for expert help in figure editing.

---

**Figure S1.**
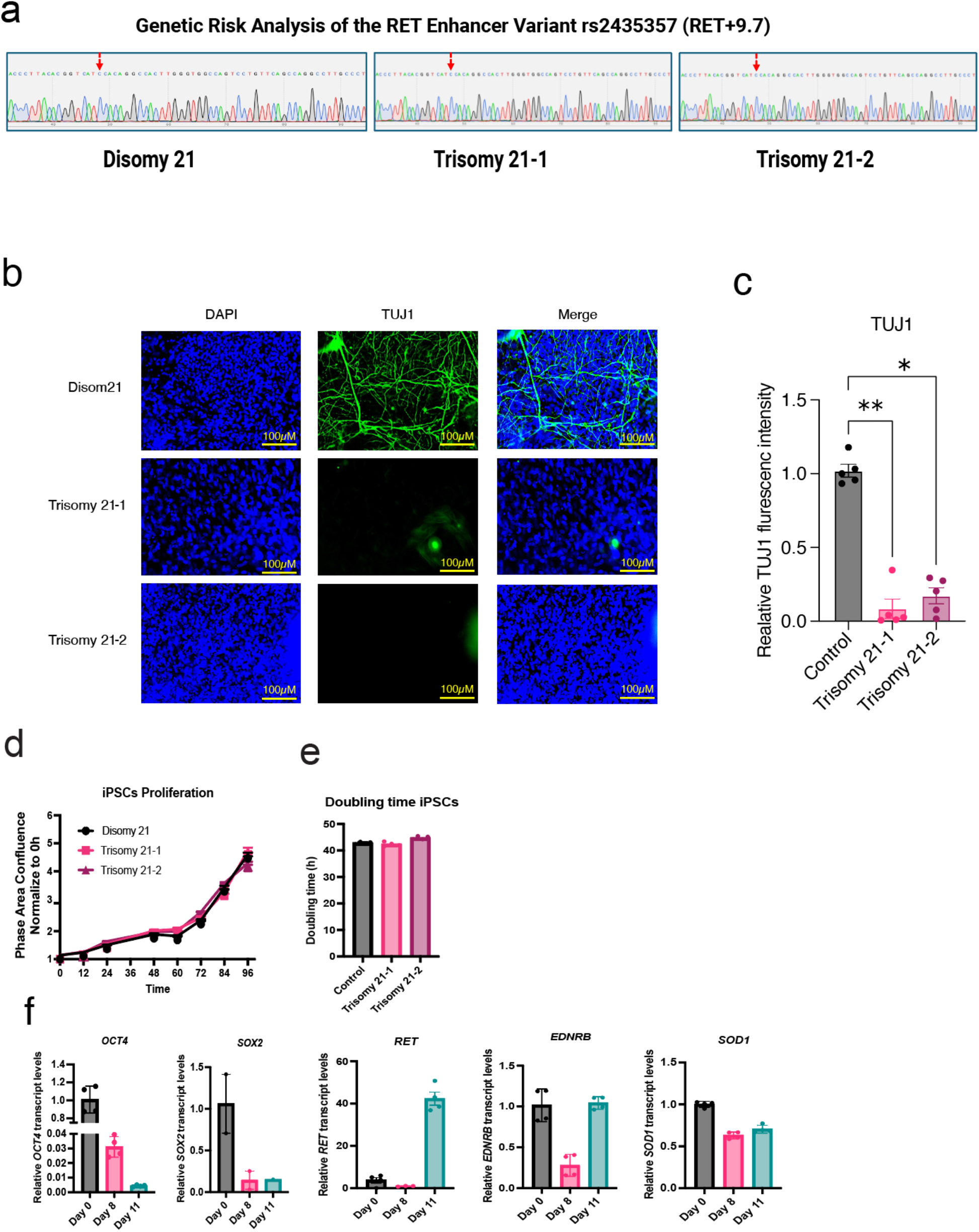
Trisomy 21 impairs neuronal differentiation of ENCDCs. **a,** Sanger sequencing of the *RET* enhancer region. A targeted genomic locus was PCR-amplified from genomic DNA of Disomy 21, Trisomy 21-1, and Trisomy 21-2 lines and sequenced to identify variant alleles within the *RET* enhancer. **b,** Immunofluorescence staining for TUJ1 in enteric neurons at Day 41 of differentiation. Disomy 21, Trisomy 21-1, and Trisomy 21-2 cells were fixed with 2% PFA and stained with an anti-TUJ1 antibody (green); nuclei were counterstained with DAPI (blue). Scale bars, 100 µm. Representative images from one of two independent experiments are shown. **c,** Quantification of TUJ1 expression per cell, computed as total fluorescence intensity normalized to total nuclei count. d, Cell proliferation assays were performed using Disomy 21 and Trisomy 21 iPSC lines. Cells were seeded at a density of 1.5 × 10⁵ cells per well in 12-well plates, and proliferation was monitored by live-cell imaging using the Incucyte system. Representative data from one of two independent experiments are shown. *n* = 3–4 wells per each group, with a minimum of 19 images analyzed per well for each iPSC line. **e,** Quantification of doubling time from cell proliferation using an R script (further details are provided in the Methods section). **f,** Relative transcript levels of *OCT4*, *SOX2*, *RET*, *EDNRB*, and *SOD1* at day 11 of differentiation in WA01 iPSCs overexpressing the indicated constructs (*n* = 2–4 biological replicates; two technical replicates per gene). Statistical significance of Fig. S1c was examined using nonparametric tests, including the Kruskal–Walli’s test with Dunn’s multiple comparisons test. Data are represented as mean ± SEM. p < 0.05, p < 0.01, p < 0.001, p < 0.0001; ns, not significant.

**Figure S2.**
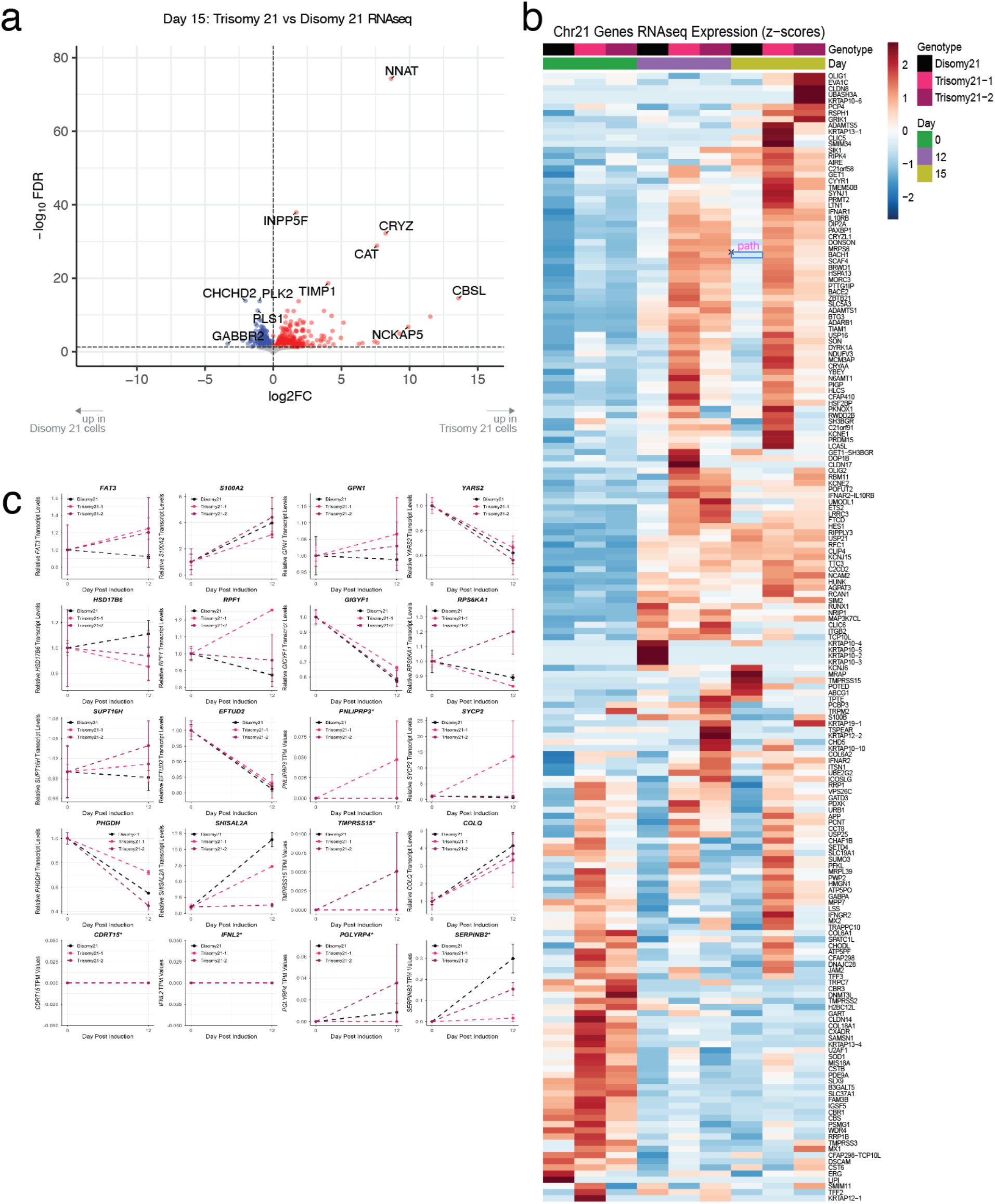
Trisomy 21 impairs HSCR genes expression. **a,** Volcano plot illustrating Trisomy 21 vs Disomy 21 significantly differentially expressed genes on day 15 post induction. **b,** Heatmap showing the expression profiles of chromosome 21 genes during differentiation. TPM expression values were z-score normalized within each gene. **c,** Relative transcript levels of novel Hirschsprung disease candidate genes, including *FAT3, S100A2, GPN1, YARS2, HSD17B6, RPF1, GIGYF1, RPS6KA1, SUPT16H, EFTUD2, PNLIPRP3, SYCP2, PHGDH, SHISAL2A, TMPRSS15, COLQ, CDRT15, IFNL2, PGLYRP4,* and *SERPINB2*.

**Supplementary Table 1.**
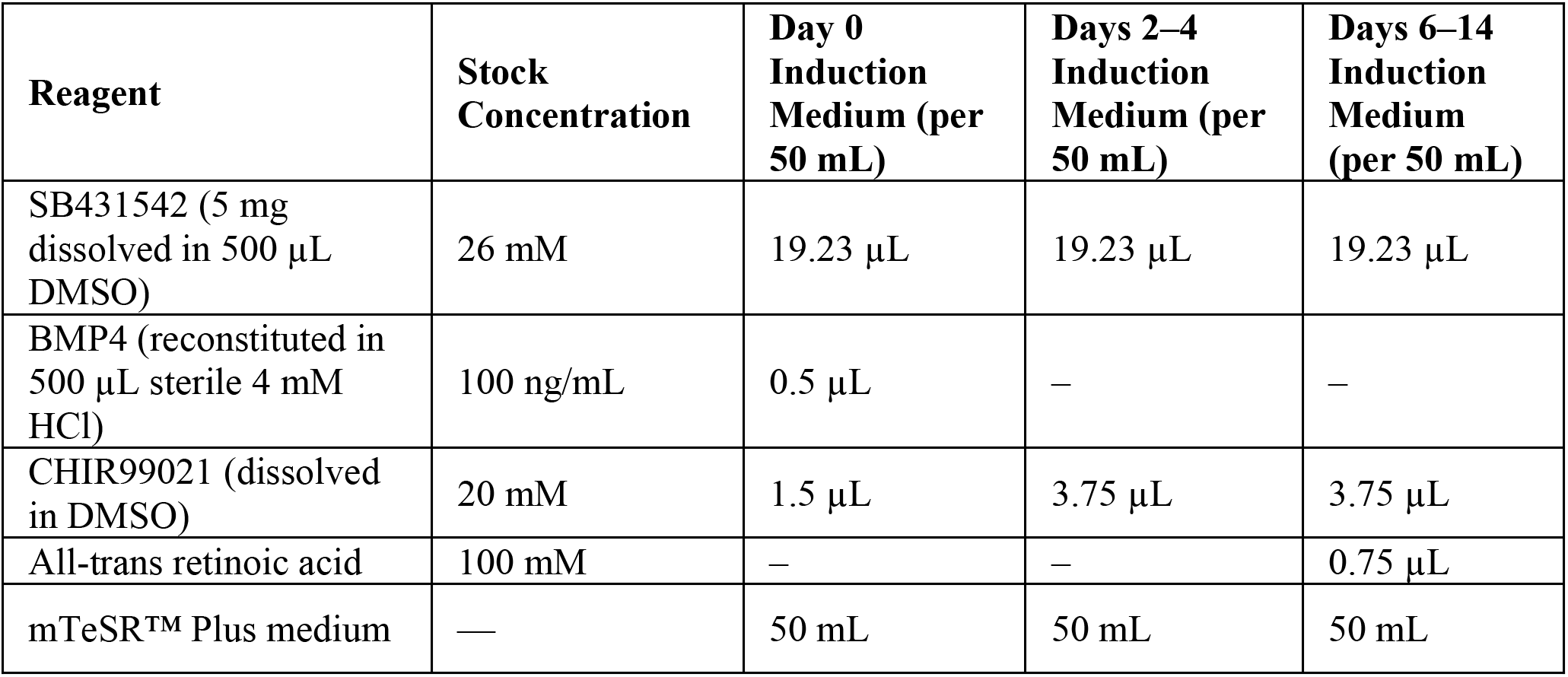

| **Reagent** | **Stock Concentration** | **Day 0 Induction Medium (per 50 mL)** | **Days 2–4 Induction Medium (per 50 mL)** | **Days 6–14 Induction Medium (per 50 mL)** |
| --- | --- | --- | --- | --- |
| SB431542 (5 mg dissolved in 500 µL DMSO) | 26 mM | 19.23 µL | 19.23 µL | 19.23 µL |
| BMP4 (reconstituted in 500 µL sterile 4 mM HCl) | 100 ng/mL | 0.5 µL | – | – |
| CHIR99021 (dissolved in DMSO) | 20 mM | 1.5 µL | 3.75 µL | 3.75 µL |
| All-trans retinoic acid | 100 mM | – | – | 0.75 µL |
| mTeSR™ Plus medium | — | 50 mL | 50 mL | 50 mL |

**Supplementary Table 2.**

| **Component** | **Final Concentration** | **Stock Concentration** | **Volume for 500 mL** |
| --- | --- | --- | --- |
| GDNF | 25 ng/mL | 125 ng/µL | 100 µL |
| Ascorbic Acid | 100 µM | 200 mM | 250 µL |
| N2 Supplement | 1× | 100× | 5 mL |
| B27 Supplement | 1× | 50× | 10 mL |
| Glutagro | 1× | 100× | 5 mL |
| MEM Non-Essential Amino Acids (NEAA) | 1× | 100× | 5 mL |
| Neurobasal Medium | — | — | 474.65 mL* |

